# No rest for the rodent: energy management strategies in the naked mole-rat

**DOI:** 10.1101/2025.11.12.688094

**Authors:** Paige Chan, Deanna Turchyn, Vincent Careau, Matthew E Pamenter

## Abstract

Organisms have access to a limited amount of energy that must be distributed among multiple physiological processes. Broadly, the total daily energy expenditure (DEE) can be partitioned into maintenance costs (i.e., resting metabolic rate RMR) and active energy expenditure (AEE). The slope (*b*) between DEE and RMR provides insights into energy management strategies. In the additive model, where changes in activity are independent of maintenance energy, DEE and RMR follow a part-whole relationship with *b*=1. In the allocation model, where increased activity requires compensatory reductions in maintenance costs, a limit on DEE causes a DEE-RMR relationship with *b*<1. In the performance model, increased activity causes an increase in maintenance costs, which causes a DEE-RMR relationship with *b*>1. Despite their high lifetime energy expenditure and resistance to age-related metabolic decline, energy management is yet to be explored in the African naked mole-rat (NMR, *Heterocephalus glaber*). To investigate metabolic strategies in the NMR, repeated metabolic and activity measurements were taken in 32 individual NMRs using a multiplexed metabolic system. DEE was not repeatable, thus the DEE-RMR covariance at the among-individual level could not be fitted. At the within-individual level, however, the positive correlation between RMR and activity and the DEE-RMR relationship with *b* > 1 indicated support for the performance model. Hence, our results indicate that within-individual changes in activity and RMR are associated, suggesting that when a NMR increases activity on a given day, the impact on DEE are disproportionate because of a concurrent increase RMR.

## Introduction

Energy is a limited currency, and every animal has an energy budget that maximizes fitness by balancing energy intake with the energy they utilize for maintenance, growth, reproduction, storage, locomotion, etc. (Bright Ross et al., 2024). Daily energy expenditure (DEE) is the total energy an organism uses in a day, which, in its simplest form, can be divided into the energy invested in basal processes (e.g., maintenance) and that invested in non-basal energy expenditure (e.g., locomotion, thermoregulation). In general, DEE increases proportionally with the incidence of energetically expensive activities such as growth (Careau, Bergeron et al., 2013), reproduction (Speakman, 2008), locomotor activity (Pagano et al., 2018; Vaanholt et al., 2007), and thermoregulation (Abreu-Vieira et al., 2015; Careau, Réale, et al., 2013; Ocobock, 2016; Vaanholt et al., 2007). By quantifying the total energy budget (i.e., DEE) and the energy invested into maintenance, one can thus infer whether compensation occurs between these two main energy budget components (Abdeen et al., 2021).

Energy invested into maintenance can be quantified through the measurement of the basal metabolic rate (BMR) of an organism. BMR represents the minimum rate of energy expenditure of a non-reproductive adult endotherm in a post-absorptive state and at its thermoneutral temperature (Lawrence, 2015). While it is desirable to eliminate as many sources of variation as possible in an experimental setting (e.g., temperature, activity, age, reproductive and prandial state), it is often impractical to meet all the BMR measurement criteria (Abdeen et al., 2021). In such cases, researchers often report resting metabolic rate (RMR), which requires only that the animal be resting, but allows for one or more of the other criteria to be violated (Speakman et al., 2004). Although only BMR truly represent an organism’s maintenance costs, RMR is typically only 5-10% higher than BMR (Nespolo et al., 2003), such that the two traits are considered analogous. Understanding the relationship between RMR and DEE can, therefore, help researchers learn how individuals balance energy commitments to obligatory functions and the facultative expenditure of energy (Bright Ross et al., 2024).

How animals manage their energy budgets can be described by one of three energy management models that predict the relationships between DEE, RMR, and activity (see Figure 1): the independent, allocation, and performance models (Abdeen et al., 2021). Specifically, if the slope (*b*) of the relationship between DEE and RMR = 1, this suggests that RMR and DEE are directly proportional to one another (i.e., a one-unit increase in RMR results in a one-unit increase in DEE). Thus, changes in locomotor activity directly impact DEE (i.e., the non-resting energy expenditure component), but do not change the maintenance. In other words, activity and maintenance are “independent”. This model is also termed the ‘additive’ model, as DEE is determined by the sum of energy invested into RMR and activity (Careau et al., 2021). A *b* < 1 indicates that allocation occurs, where additional energy expenditure associated with locomotor activity must be compensated for with a decrease in RMR, and *vice versa* (Mathot & Dingemanse, 2015). This is also termed the ‘compensation’ or the ‘constrained’ model, as energy budgets are constrained (Pontzer et al., 2015), which forces a negative relationship between RMR and locomotor activity. Finally, when increases in activity increase the maintenance energy cost, as reflected in RMR, the DEE-RMR relationship will have a *b* > 1. This “performance” model proposes that the cost of behaviour goes beyond the energy required to express the behaviour itself, because sustaining high levels of activity requires the maintenance of a larger metabolic machinery (Daan et al., 1990).

**Figure 1.**
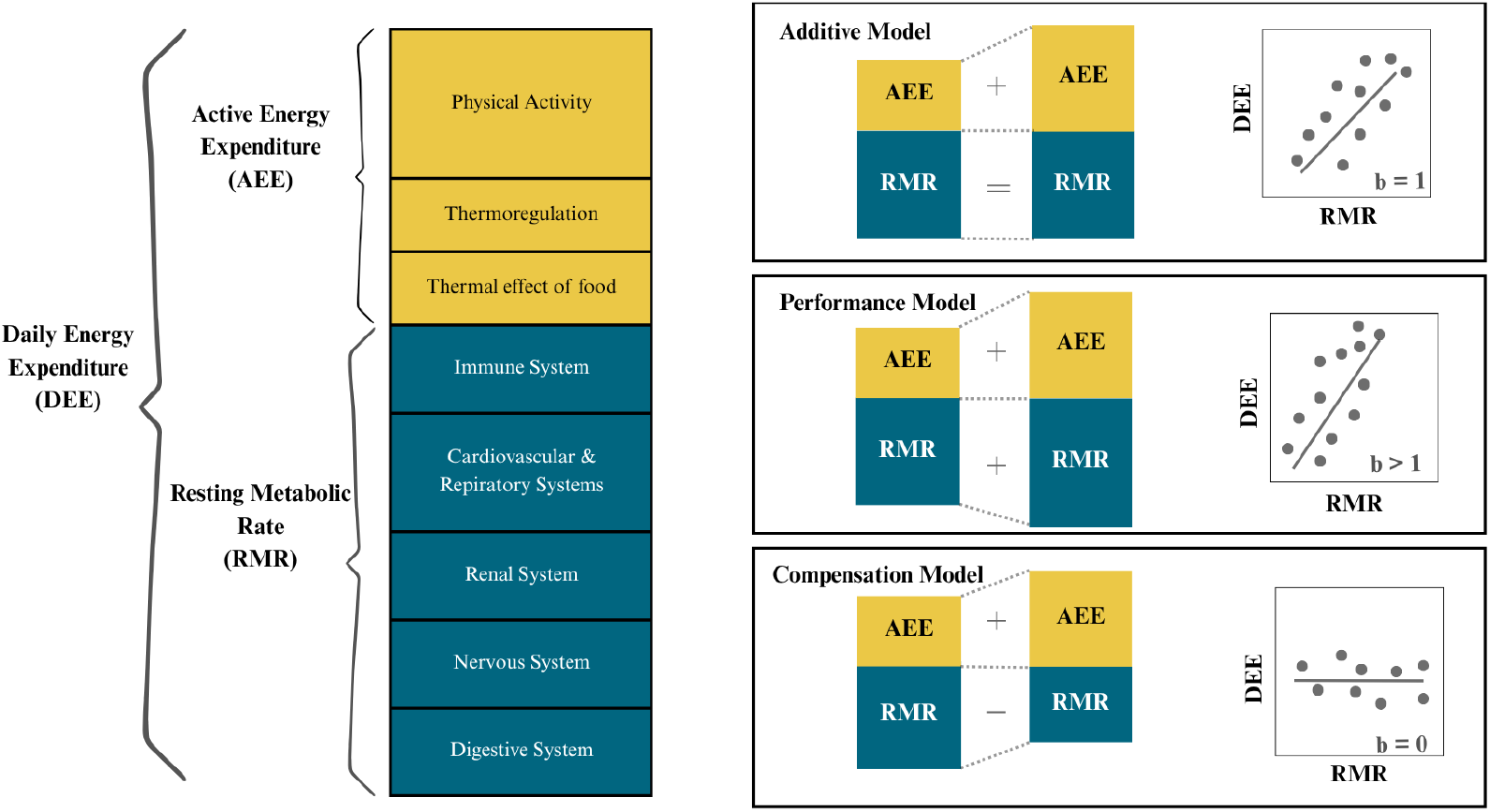
Conceptual framework of energy budgeting and energy management models in naked mole-rats (*Heterocephalus glaber*). The components of daily energy expenditure (DEE) in adult, non-reproductive and thermoneutral animals. Energy management models illustrating different relationships between resting metabolic rate (RMR) and DEE. Modified from Careau et al. (2021).

Studying the relationship between RMR and activity requires the covariance of these traits to be partitioned at the among- and within-individual levels, as they are both versatile traits (Dingemanse & Dochtermann, 2013). Which energy management model applies among- and within-individuals can be determined by partitioning the relationship between RMR and DEE at each level using repeated measurements from the same individuals (Careau et al., 2019). Partitioning covariances at each level is critical as patterns among individuals might hide important processes occurring within individuals, especially if different models apply at each level and cancel each other. For example, Abdeen et al. (2021) partitioned the covariances in DEE and RMR at the among- and within-individual levels in white-footed mice (*Peromyscus leucopus)* using repeated measurements and discovered that the results supported the performance model at the among-individual level, but at the within-individual level, the allocation model was supported.

One animal that is of particular interest in considering energy management models are African naked mole-rats (NMRs; *Heterocephalus glaber*). NMRs have lower metabolic rates than other eutherians of the same size, which do not tend to change as they age, causing them to have a lifetime energy expenditure 4 times greater than that of mice, despite low BMR (Riccio and Goldman, 2000; Yap et al., 2022; O’Connor et al., 2002). NMRs also have a very plastic metabolism and can maintain consciousness and activity when their metabolic demand is decreased by > 85% in environmental hypoxia. Beyond this flexible metabolism, most members of a given NMR colony are sexually supressed subordinate animals that do not go through puberty (Bennett et al., 2022). Thus, NMRs are unique among mammals regarding energetic trade-offs because most adults do not invest energy into reproduction. Finally, NMRs are subterranean and life in underground burrows can be energetically expensive, especially for worker animals, as their food resources, consisting of tubers and roots, are more dispersed and are therefore energetically costly to obtain (Vleck, 1979). How NMRs remain active despite metabolic flexibility is thus an intriguing question of energy budgeting.

The main objective of this study was to describe how NMRs distribute energy between maintenance versus behavioural activities by quantifying the relationships between RMR, mass, home cage activity and DEE. Repeated measurements of locomotor activity, DEE, RMR, and body mass were collected from non-breeding NMRs to estimate the slope (*b*) between DEE and RMR at the among- and within-individual levels, while also investigating the relationships between home-cage activity, mass, DEE and RMR.

## Materials and Methods

### Study animals

NMRs were housed in colonies in multi-cage systems at 30ºC with 21% oxygen, 50% humidity and a 12L:12D light cycle. A total of 32 non-breeding animals were randomly selected and divided into 4 groups of 8. All procedures conducted during this experiment were approved by the University of Ottawa Animal Care Committee (protocol: #4415).

### Experimental design

This study replicates the approach of Abdeen et al. (2021), which was based on repeated 24-h metabolic and behavioural measurements. Initial body mass measurements were collected from each animal before the start of the experiment, with an average mass of 50.5 ± 7.24 g. Individual NMRs were randomly placed into one of eight metabolic chambers of an 8-channel multiplexed Promethion System (Sable Systems International, North Las Vegas, USA), and left undisturbed for 24 h in an enclosed and sound-dampened room held at 30ºC, with 21% oxygen, 50% humidity and a 12L:12D light cycle. The metabolic cages contained food, consisting of bedding, raw sweet potato and rodent chow soaked in water. No water was provided to the animals because NMRs acquire their water through food intake (Jarvis & Sherman, 2002).

The Promethion system is an open-flow multichannel respirometry system that measures O_2_ consumption, CO_2_ production, water vapour loss, and locomotor activity. Locomotor activity was continuously measured within each cage using the BXZ-1 Beambreak Activity Monitor. Incurrent and excurrent gases and water vapour levels were measured and multiplexed across all 8 cages by the Promethion GA-3 Gas-Analyzer through the fuel cell O_2_ analyzer, spectrophotometric CO_2_ analyzer, and the capacitive water vapour partial pressure analyzer, respectively. Raw O_2_ consumption and CO_2_ production traces collected from the Promethion System were used to calculate metabolic rate using the Weir equation (1949). The lowest 30 min period of metabolic rate (measured using six 30s measurements collected at 5-minute intervals), without any locomotor activity, was used to calculate RMR. The entire 24 h period of metabolic measurement was averaged to calculate DEE.

### Statistical analysis

Parameters of interest were estimated using a 4-trait multivariate mixed model fitted in ASReml-R 4.0 (Butler et al. 2018). The dependent variables were DEE, RMR, body mass, and locomotor activity. Home-cage activity (in m·h^-1^) was square-root transformed to improve the normality of the residuals. DEE and RMR were kept in their original scale (kcal/hr). The fixed effects within this model included test chamber (chambers 1-8) and test sequence and were fitted separately for each of the four traits. Individual identity was included as a random effect to partition the phenotypic variance (*V*_P_) of each trait into variance among individuals (*V*_among_) and within individuals (*V*_within_; residuals). Individual repeatability (*R*) was calculated using the equation *R* = *V*_among_/(*V*_among_ +*V*_within_). Since DEE and RMR were not repeatable (see Results), there were no among-individual differences in RMR and DEE and a model specifying covariances at the among-individual level did not converge. Therefore, covariances were fitted only at the within-individual (residuals) level. Correlations at the within-individual level were calculated by dividing the covariance by the square root of the product of their variances.

The slope (*b*) between DEE and RMR was calculated using the (co)variance estimate from the multivariate fixed model to determine which energy management model applied to NMRs at the within-individual level. Calculating the slope between DEE and RMR describes the relationship between the resting and non-resting constituents of the NMR energy budget. The within-individual DEE-RMR slope (*b*_within_) was calculated using two regression methods, the ordinary least squared (OLS) and standard major axis (SMA) methods, which can differ depending on the ratio of measurement error in DEE to RMR (Halsey & Perna, 2019). The OLS slope was calculated by dividing the DEE-RMR covariance by the RMR variance, and the SMA slope was calculated by taking the square root of the ratio of the variance of DEE over the variance of RMR. The standard error (SE) estimates for the slope were calculated using the delta method.

## Results

### Repeatability of DEE, RMR, body mass, and activity

Repeatability analyses indicated that DEE was not repeatable (Table 1). RMR showed very low repeatability (*R* = 0.08 ± 0.04; Table 1). In contrast, body mass exhibited high repeatability (*R* = 0.95 ± 0.02) and home-cage activity was moderately repeatable (*R* = 0.38 ± 0.12). Correlations estimated at the within-individual level indicated that activity was positively correlated with both RMR (*r* = 0.51 ± 0.04) and DEE (*r* = 0.54 ± 0.08; Table 1). The DEE-RMR *b*_within_ estimates exceeded 1. Using ordinary least squares, the within-individual slope (b_within_) was 2.04 ± 0.12, while standardized major axis regression produced a slope of 1.89 ± 0.12 (Table 1; Figure 2).

**Table 1.** Summary of key effect sizes (±se) from a multivariate mixed model analysis of daily energy expenditure (DEE; kcal/hr), resting metabolic rate (RMR; kcal/hr), body mass (g), and home-cage activity (m/hr) in 32 subordinate naked mole-rats (*Heterocephalus glaber*). Values represent repeatability estimates, trait correlations at the within-individual level and slope estimates to assess support for energy management models. Ordinary least squared (OLS) and standard major axis (SMA) slopes were both calculated at the within-individual level.

| Analysis Type | Trait(s) Analyzed | Result | Interpretation |
| --- | --- | --- | --- |
| Repeatability | DEE | $0 \pm 0.06$ | Not repeatable |
| Repeatability | RMR | $0.08 \pm 0.04$ | Not repeatable |
| Repeatability | Activity | $0.38 \pm 0.12$ | Moderately Repeatable |
| Repeatability | Mass | $0.95 \pm 0.02$ | Highly repeatable |
| Correlation | RMR vs. Activity | $0.51 \pm 0.04$ | Positive correlation |
| Correlation | Activity vs. DEE | $0.54 \pm 0.08$ | Positive correlation |
| OLS slope ($b_{\text{within}}$) | RMR vs. DEE | $2.04 \pm 0.12$ | Performance model |
| SMA slope ($b_{\text{within}}$) | RMR vs. DEE | $1.89 \pm 0.12$ | Performance model |

**Figure 2.**
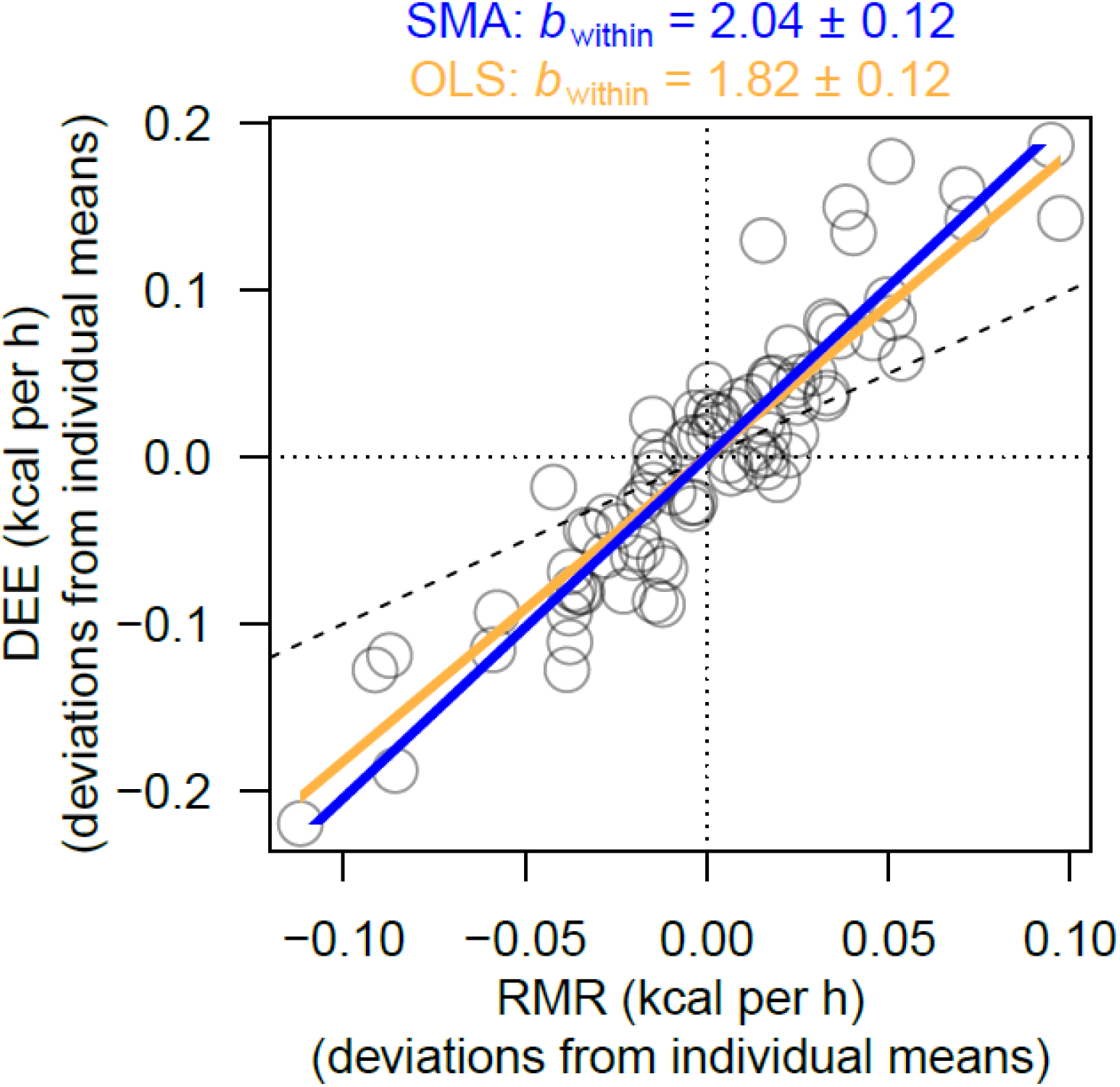
Representation of the within-individual slopes (*b*_within_) between daily energy expenditure (DEE; kcal/hr) and resting metabolic rate (RMR; kcal/hr) in 32 individual naked mole-rats (*Heterocephalus glaber*). Ordinary least squared (OLS; in yellow) and standard major axis (SMA; in blue) slopes are presented in both cases. Dashed line represents the 1:1 identity line, where y = x.

## Discussion

This study is the first to describe energy management and the relationships between RMR, DEE, body mass, and activity in NMRs. By looking at the DEE-RMR slopes, we found that the performance model was supported at the within-individual level, as seen with the OLS and SMA slopes being greater than 1. Accordingly, activity was positively correlated with both RMR and DEE at the within-individual level. The DEE-RMR slopes and correlations could only be partitioned at the within-individual level because DEE was not repeatable (and repeatability in RMR was very low, while activity was moderately repeatable and body mass was highly repeatable).

The lack of repeatability in DEE is unexpected yet provides an interesting avenue for interpretation. There are a few explanations for why NMRs may exhibit repeatability values of zero for DEE and near zero for RMR. First, high within-individual variation across trials can mask any among-individual differences, effectively reducing repeatability to zero. One potential source of this variation could be stress induced by reintroducing animals into their home colonies after each trial. Upon reintroduction, subjects were often followed, surrounded and nipped at by cage mates, triggering a temporary frenzy within the colony. In some cases, returning subjects even led to aggressive and violent fights involving multiple animals. This social upheaval could have caused a conditioned stress response in the animals, whereby the act of transporting an animal from their colony to the metabolic cages in the laboratory became associated with anxiety. As a result, subjects may have exhibited altered physiological states during later trials, creating more within-individual variation. Thus, RMR and DEE might not be traits with strong, stable differences in NMRs. Second, it is theoretically possible that there is no true among-individual variation in RMR or DEE. However, this is unlikely, as these are complex traits influenced by size, organ function and personality traits, all of which are different among individuals. Finally, experimental design and/or measurement error could have influenced the repeatability of RMR and DEE. Having a small number of repeated measurements per individual can create a considerable amount of measurement noise, reducing the ability to detect individual variation among animals.

Although RMR and DEE had very low repeatability, body mass and home-cage activity were significantly repeatable. One may ask ‘if activity is a component of DEE, then why is DEE not repeatable when activity is?’ A possible explanation to this is that RMR contributes more to DEE than activity. As mentioned previously, RMR can largely vary within an individual between trials and since RMR contributes more to DEE, this can mask the variation of activity, ultimately resulting in DEE not being repeatable. Therefore, the repeatability of body mass and activity suggests stable individual differences, whereas the lack of repeatability in DEE and RMR suggests that these traits may be more plastic and influenced by environmental and internal factors, such as stress or social dynamics.

At the within-individual level, RMR and home-cage activity were positively correlated, as well as home-cage activity and DEE (0.51 ± 0.04 and 0.54 ± 0.08) respectively. These results suggest that energy investment in maintenance metabolic functions increases with home-cage activity, or vice versa, and that DEE increases as home-cage activity increases, indicating that home-cage activity contributes positively to DEE. This supports the performance model, indicating that NMRs may upregulate overall energy output rather than compensating between maintenance functions and activity. This is consistent with the results of the DEE-RMR slopes that also indicate support for the performance model at the within-individual level. Having *b*_within_ > 1 reinforces the idea that DEE increases more than expected as RMR increases. It is therefore suggested that NMRs pay an extra maintenance cost related to an increased RMR in addition to the high energy costs related to their activity.

Interestingly, Abdeen et al. (2021) discovered that white-footed (*Peromyscus leucopus)* mice showed support for the performance model at the among-individual level and the allocation model at the within-individual level (OLS *b*_among_ = 1.57; SMA *b*_among_ = 2.66; OLS *b*_within_ = 0.47; and SMA *b*_within_ = 0.83). Although we were unable to determine which energy management model NMRs support at the among-individual level due to the lack of repeatability in DEE, we discovered that NMRs support the performance model at the within-individual level. The difference in which energy management model is supported at the within-individual level between both species clearly indicates that there are differences in how NMRs and white-footed mice manage their energy budgets. Looking at the slope of the DEE-RMR relationship at the within individual level in a naked mole-rat, every unit increase in RMR results in a 2 unit increase in DEE, compared to the white-footed mouse whose DEE remains nearly the same per unit of increase in RMR. These differences are likely due to species-specific ecology and behaviour, evolutionary history, and social structure. For example, white-footed mice are solitary animals that live in environments with higher predation pressure, so they must juggle reproduction, survival, and foraging on their own (Aguilar, 2011). As their environments are unpredictable, adopting a re-allocation strategy may be advantageous for mice where they can switch between conserving and spending energy, and can make trade-offs between RMR and active energy expenditure. On the other hand, as NMRs are eusocial, they have more fixed roles where labour is divided within the colony. Their energy systems may thus be shaped for long-term specialization in stable environments, where the performance model is consistently supported so that higher RMR can fuel sustained activities. However, this interpretation is highly speculative at this point (Garland & Adolph, 1994), and there exists many other factors that can result in different energy management strategies between NMRs and white-footed mice (i.e., such as the presence or absence of fur).

Increasing RMR to support high active energy expenditure may be advantageous for NMRs as their bodies can bear their high-functioning lifestyles. Having more energy invested into maintenance functions, such as the cardiovascular and respiratory systems, allows for sustained effort in their daily tasks, such as foraging, burrowing, and alloparental care. In addition, having more energy allocated towards maintenance systems can have a crucial role in the NMRs known resilience to aging and disease. NMRs can live for up to 30 years, which is largely attributed to sustained good health and cancer resistance, suggesting they have strong cellular repair and immune function (Edrey et al., 2011). An assumption in ecology is that immune responses are costly, where their costs must be reflected in a need to increase metabolic rate or trade-off with other life processes, such as reproduction or growth (Brace et al., 2018). Therefore, support of the performance model at the within-individual level may be a reason to why NMRs are resistant to age-related disease and cancer, as they can flexibly allocate more energy into activity and maintenance functions, such as immunity and potentially DNA repair systems.

In our study, we use non-breeders, consisting of both “subordinates” males and females that are ranked lower than the breeder males and the queen. Interestingly, non-breeders form dominance hierarchies among themselves, in which their rank is highly correlated with their body weight and can be observed by which animal walks over the other when crossing tunnels (Clarke & Faulkes, 1997; Clarke & Faulkes 1998; Toor et al., 2015; Edwards et al., 2020). Furthermore, non-breeders are divided into workers, soldiers, and dispersers. First, workers are typically highly interactive with other colony members and spend most of their time working or foraging, huddling in the nest, or contributing to alloparental care, such as carrying pups (Holmes et al., 2021). In addition, research suggests that worker productivity may be increased depending on the queen’s aggression, as she exhibits more shoving behaviour toward ‘lazy’ workers to get them to work harder (Reeve, 1992). Next, soldiers are the least interested in the pups and are the most aggressive of the non-breeders, specifically toward unfamiliar animals, as they act as the first line of defence to protect their colony (Lacey et al., 1991). Finally, although this role is questioned in the field, the dispersers try to leave their colony or explore by digging burrows, where the motivation behind leaving the colony may be sex-specific (Holmes et al., 2021). Therefore, these distinct roles within the colony highlight the varying activity levels and energetic demands required for each individual. Thus, a clear limitation of our study arises from the lack of information on the castes within the NMR hierarchy, which may have obfuscated among- and within-individual individual variation depending on the stability versus changes in status. Future studies should determine the specific castes of measured animals, which may offer insights into our understanding of NMR energy management

## Conclusions

We investigated the relationships between RMR, DEE, and home-cage activity in subordinate NMRs. Our findings revealed that while body mass and activity were significantly repeatable, indicating consistent individual differences, DEE was not repeatable and repeatability in RMR was very low. This suggests that metabolic traits in NMRs are highly plastic and may be strongly influenced by internal or environmental factors, such as stress or social interactions. Due to the lack of repeatability in DEE, among-individual covariance with this trait could not be assessed. However, at the within-individual level, there was a positive correlation between RMR and activity, and the DEE-RMR relationship had a *b* > 1, both indicating support for the performance model. This suggests that an individual who expend more on energy than usual on activity also invest more energy than usual in maintenance functions, resulting in a disproportionate increase in overall DEE. Overall, these results highlight the importance of within-individual flexibility in energy budgeting and suggest that NMRs rely on dynamic physiological adjustments, which may be advantageous in context of their cooperative and high-functioning lifestyles.

Overall, these findings enhance our understanding on energy management strategies in NMRs and adds to the existing theoretical models of animal energetics. This work establishes the first quantitative assessment of energy management in the naked mole-rat and provides a basis for future research into how their unique physiology supports long lifespan and metabolic stability.

## Abbreviations

DEE: daily energy expenditure
NMR: naked mole-rat
OLS: ordinary least squares
RMR: resting metabolic rate

